# Reliable detection and quantification of plasmodesmal callose in *Nicotiana benthamiana* leaves during defense responses

**DOI:** 10.1101/2023.09.30.560305

**Authors:** Amie F. Sankoh, Joseph Adjei, Daniel M. Roberts, Tessa M. Burch-Smith

## Abstract

Callose, a beta-(1,3)-D-glucan polymer, is essential for regulating intercellular trafficking via plasmodesmata (PD). Pathogens manipulate PD-localized proteins to enable intercellular trafficking by removing callose at PD, or conversely by increasing callose accumulation at PD to limit intercellular trafficking during infection. Plant defense hormones like salicylic acid regulate PD-localized proteins to control PD and intercellular trafficking during innate immune defense responses such as systemic acquired resistance. Measuring callose deposition at PD in plants has therefore emerged as a popular parameter for assessing the intercellular trafficking activity during plant immunity. Despite the popularity of this metric there is no standard for how these measurements should be made. In this study, three commonly used methods for identifying and quantifying PD callose by aniline blue staining were evaluated to determine the most effective in the *Nicotiana benthamiana* leaf model. The results reveal that the most reliable method used aniline blue staining and fluorescent microscopy to measure callose deposition in fixed tissue. Manual or semi-automated workflows for image analysis were also compared and found to produce similar results although the semi-automated workflow produced a wider distribution of data points.

## Main text

In plants, cell-to-cell communication is largely mediated by numerous small pores in the cell wall called plasmodesmata (PD) that directly connect the cytoplasms of neighboring cells (Kang et al., 2022). PD are essential for intercellular communication, plant development, and growth. In many instances PD trafficking is regulated by the controlled accumulation of the β-1,3-glucan, callose, in the cell wall surrounding plasmodesmal pores (Zavaliev et al., 2011; Sankoh and Burch-Smith, 2021). Callose deposition restricts intercellular trafficking, whereas callose degradation increases intracellular trafficking. The restriction of PD trafficking likely occurs during defense against pathogens through the physical closure of the PD pore by the accumulation of callose (Zavaliev et al., 2011; Singh et al., 2017).

Plant hormones regulate PD callose levels to adjust intercellular trafficking. Exogenous application of salicylic acid (SA) to plants activates immune responses with a concomitant induction of callose accumulation at PD that lowers PD-mediated intercellular trafficking. SA induces the activity of PLASMODESMATA LOCATED PROTEIN (PDLP)5-directed machinery that induces callose accumulation and close PD during innate immune responses (Lee and Lu, 2011; Lee et al., 2011). Local pathogen infection often leads to resistance in systemic healthy, uninfected tissues by a process termed systemic acquired resistance (SAR). Systemic transport of SA is important for SAR, although it is transported in the apoplast and not via intercellular transport through PD (Kachroo and Kachroo, 2020). In contrast to SA, other SAR signaling molecules azelaic acid (AzA) and glycerol-3-phosphate (G3P) move systemically via PD (Lim et al., 2016). The importance of PD in SAR was highlighted by the observation of defective SAR in *pdlp1* and *pdlp5* mutants (Lim et al., 2016). This is proposed to possibly result from altered PD permeability due to altered callose metabolism (Lee et al., 2011; Caillaud et al., 2014). SAR is also linked with the enhanced levels of callose accumulation upon secondary pathogen inoculation (Lee and Hwang, 2005; Conrath, 2006). Thus, many of SA’s roles in innate immunity may link to role in regulating PD via callose metabolism.

Given the importance of callose to regulating PD trafficking, it has become common practice to measure PD callose levels, for example, in response to pathogen infection as a gauge of the plant’s response to infection. Because of its ability to bind callose (β-1,3-glucan), cellulose, and related polysaccharides in the cell walls, aniline blue has been widely adopted for visualization of callose deposition at PD (recent examples include (Huang et al., 2022; Muller et al., 2022; Yan, 2022)). We noticed that procedures for measuring PD callose varied between labs, with important differences in how samples were treated before application of aniline blue. In several studies, samples were fixed before staining (e.g. (Lee and Hwang, 2005; Lee et al., 2011; Zavaliev and Epel, 2015)), while in others aniline blue was introduced into living tissues without fixation (e.g. (Caillaud et al., 2014; Muller et al., 2022; Rocher et al., 2022). Because of these disparate protocols, potential issues arise when comparing studies that use different staining approaches. The purpose of the present study was to determine which aniline blue staining approach was most reliable when used with leaves from *Nicotiana benthamiana*, a widely used model for plant-pathogen interactions (Goodin et al., 2008). For this purpose, three representative protocols were tested to identify the most reliable protocol for callose staining with aniline blue in *N. benthamiana* leaves.

The LAB cultivar of *N. benthamiana* was used in this study (Naim et al., 2012; Bally et al., 2015). Seedlings were germinated on soil for 7-10 days before transplanting to individual pots and plants were grown on light carts under long day conditions with 16 hours light (120 μmol m^-2^ s^-1^) and 8 hours dark at 25 °C. Miracle Gro All-purpose Plant Food fertilizer was applied after 10-13 days post germination (3 days after transplanting). The leaves of 4-5 weeks old plants were used for callose quantification and salicylic acid-treatment experiments.

The first protocol for callose staining we tested, Method 1, was adapted from (Zavaliev et al., 2011). An entire cut leaf was submerged in 95-96% ethanol in a 500-mL polypropylene jar to fix and bleach the tissue. The petiole was held with forceps to avoid mechanical damage to the leaf. The jar was sealed, and samples were incubated at room temperature on a shaker at 30-40 rpm for at least 5 hours. The ethanol solution was changed after 2 hours incubation to accelerate tissue bleaching. Incubation was continued until the leaves were completely or nearly completely bleached, but less than 6 hours, since plasmolysis occured after this point. Bleached leaves were removed and were placed in a petri dish and cut into 5 mm wide strips with a razor blade. The cut strips were rehydrated in double distilled water (DDW) with 0.01% (v/v) Tween-20 and incubated at room temperature for 1 hour on a shaker at 30-40 rpm. Tissues were then transferred to a small petri dish (35mm in diameter and 10 mm height) and submerged in 1% (w/v) aniline blue (Ward’s Science plus, Rochester NY LOT AD-22103) in 0.01 M K_3_PO_4_, pH 12. The uncovered petri dish was placed in a desiccator under house vacuum for approximately 10 minutes followed by slow release of pressure. The petri dish was covered and sealed with foil before shaking at room temperature for 2 hours at 30-40 rpm. The samples were then directly imaged without destaining with a Leica SP8 laser scanning confocal scanning microscope (Leica, Whetzlar, Germany).

Callose deposits in the abaxial side of the leaves were imaged using a 40x/1.10 water immersion objective (HC PL APO CS2 40x/1.10 WATER). The excitation for aniline blue was carried out with a 405-nm laser, UV (0.5-1%) and emission was captured from 415 nm-525 nm. Z-stacks were collected from multiple regions of interest (ROI; approximately 23-25 ROIs per leaf) using the system optimized step size (<1 μm). To observe as many PD as possible, the pinhole aperture was initially set to open. Laser intensity and gain were adjusted to optimize the PD fluorescence against the background signal, while being careful to avoid oversaturation of callose sites. Saturation of the stomata was permitted. Twenty three to twenty five Z-stacks were imaged using the same settings of image acquisition in the independent experiments.

Because individual PD cannot be resolved by light microscopy, we measured the clusters of callose at PD in pitfields, which are concentrated areas of PD between 0.05 μm^2^ and 4 μm^2^ in size. Representative images obtained in maximum intensity z-stack projections from confocal microscopy of the untreated control (Fig. 1A) and SA treatment (Fig. 1B) are shown. The control shows multiple pitfields in the cell wall, and other artifacts that are not within the cell wall (marked with white arrows).

**Figure 1.**
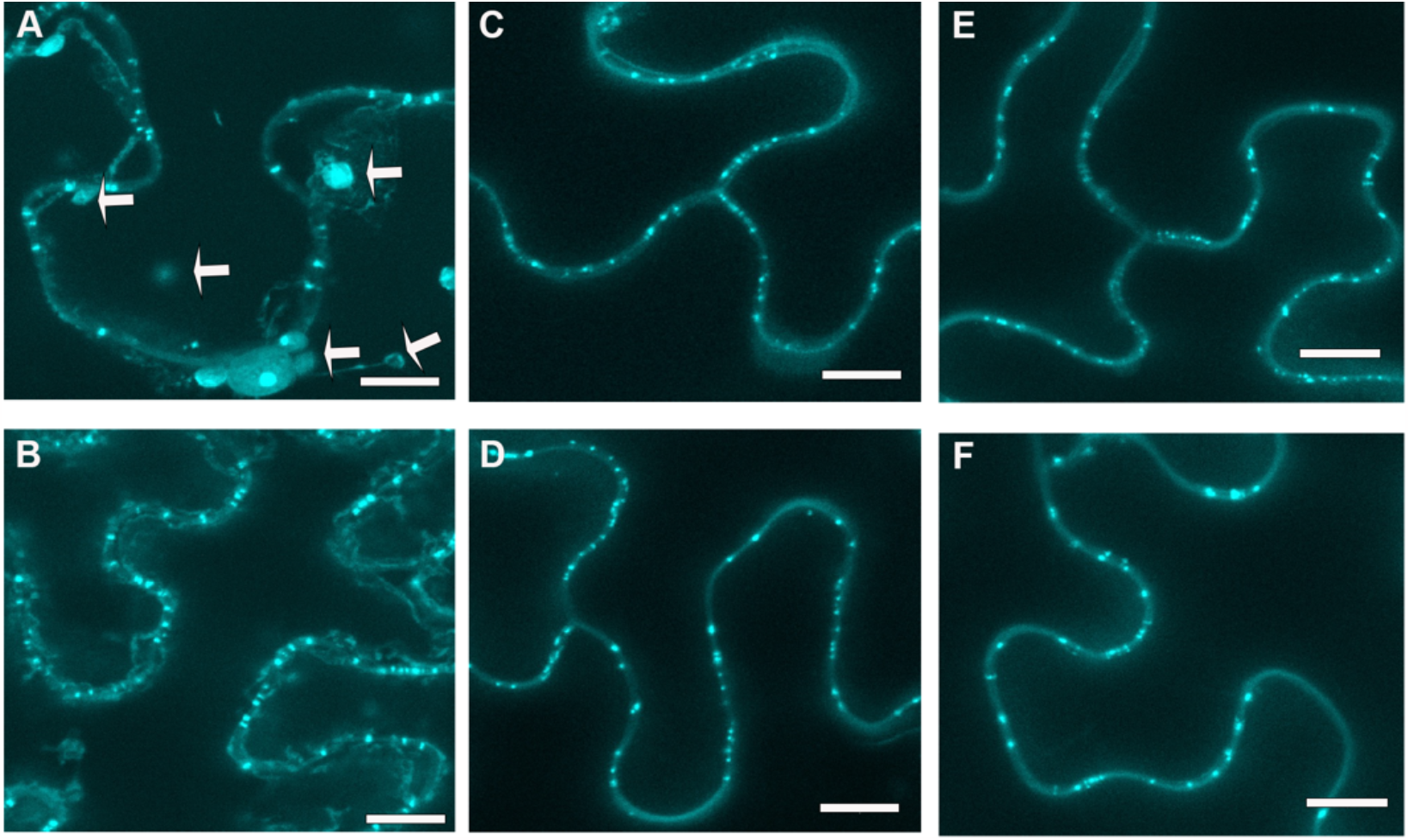
Results from the three methods of aniline blue staining for plasmodesmata pitfield imaging. *N. benthamiana* leaf epidermal cells with staining of plasmodesmata (PD)-associated callose using 1% Aniline blue concentration. Representative images are shown for three different callose staining methods, both for untreated and SA treated plants. Panels corresponding to the three methods of aniline blue staining. **A**, Method 1 using fixed tissue. **B**, Method 1 in tissue treated with 0.5 mM SA for 12 hours. **C**, Method 2 using non-fixed tissue directly infiltrated with aniline blue. **D**, Method 2 in SA-treated tissues. **E**, Method 3 using non-fixed tissue stained with aniline blue for 30-60 minutes. **F**, Method 3 in SA-treated tissues. Images are maximum projections of Z-stacks of images collected by confocal fluorescence microscopy. Scale bars = 10μm.

The other methods employed involved the direct staining of plant tissue without a prior fixation step. The aniline blue staining solution and confocal microscopy imaging protocols were the same as those used for Method 1. Method 2 was adapted from a protocol available at https://www.jic.ac.uk/research-impact/aniline-blue-staining-of-qunatification-of-plasmodesmal-callose/. Staining involved direct infiltration of 1% (w/v) aniline blue into leaves while they were still attached to the plants using a 1-mL needleless syringe. A representative maximum intensity projection of a z-stack is shown with and without SA treatment (Fig. 1C and D). Aniline blue foci representing PD were clearly observed in the cell wall, and staining of other structures were rarely observed.

Method 3 was adapted from (Cui and Lee, 2016). Leaves were cut into 5mm strips and incubated in 1% (w/v) aniline blue in 0.01 M K_3_PO_4_, pH 12 in darkness for 30-60 minutes. Representative images with and without SA treatment are shown (Fig. 1E and F). Similar to Method 2, aniline blue-stained foci were only detected in the cell wall, and there were few, if any, other cellular structures were stained (Fig. 1).

To determine the average density of PD pitfields, at least 25 z-sacks were collected from three to four biological replicates for each method, and analysis was performed using ImageJ Fiji (Schindelin et al., 2012). Pitfields were quantified per 100 μm^2^ of cell wall for a region of interest. The workflow of the analysis for identification of callose deposition at PD is shown (Fig. S1 using the data presented in Fig. 1A). The first step in the image analysis workflow was to convert the maximum intensity projection to 8-bit (Fig. S1). The image was then inverted to perform inverse binary thresholding to measure the area of pitfields shown in red spots (Fig. S1). Cell-wall lengths were measured by tracing the cell wall in the region of interest (Fig. S1). Only the total number of pitfields within the traced cell wall was calculated within each region of interest as pitfields/unit area (Fig. S1).

Salicylic acid can reduce intercellular trafficking via PD during innate immune response against pathogens (Koo et al., 2020). To compare the various methods for the quantitation of the accumulation of callose at PD induced by SA, *N. benthamiana* leaves were treated with 0.05 mM SA, and after 12 hours the leaves were stained with aniline blue using each of the three previously described methods. We analyzed multiple datasets of images from SA-treated *N. benthamiana* plants along with control plants. A comparison of the PD density in control and SA-treated plants from multiple experiments is shown in Fig. 2. The colors represent data points from various biological replicates. An interquartile range (IQR) analysis was used to identify and exclude outlier data points more than 1.5 times the interquartile range (Fig. 2B). All three methods display similar medians in the control plants. However, the three methods showed distinct sensitivity and reproducibility/consistency when comparing SA-induced aniline blue stained PD foci (Fig. 2). For Method 1, each replicate showed an increase in PD callose, reflected as an increase in PD density, with median pitfield density increasing from 3.4 to 5.7 on treatment with SA (p<0.0005; Fig. 3A). For method 2, however, the experimental replicates did not consistently show increased callose at PD in response to SA treatment (Fig. 3B). This incosistency is best exemplified by Rep 2 (green dots) and Rep 4 (black dots), which show no change or a decrease in pitfield density upon SA application, respectively. It is worth noting that applying IQR analysis to the datasets resulted in a change in the statictical significance of the results obtained with method 2 going, with no significant difference before IQR analysis compared to statistically significant difference after elimination of outlying data points by IQR analysis (compare Fig. S2 to Fig. 3). Method 3 showed a similar inconsistency with significant differences between the control and SA treated plants only observed for one biological replicate (Rep 2 in Fig. 3C). The median pitfield density increased from 3.2 to 4.6 on treatment with SA (p<0.0005).

**Figure 2.**
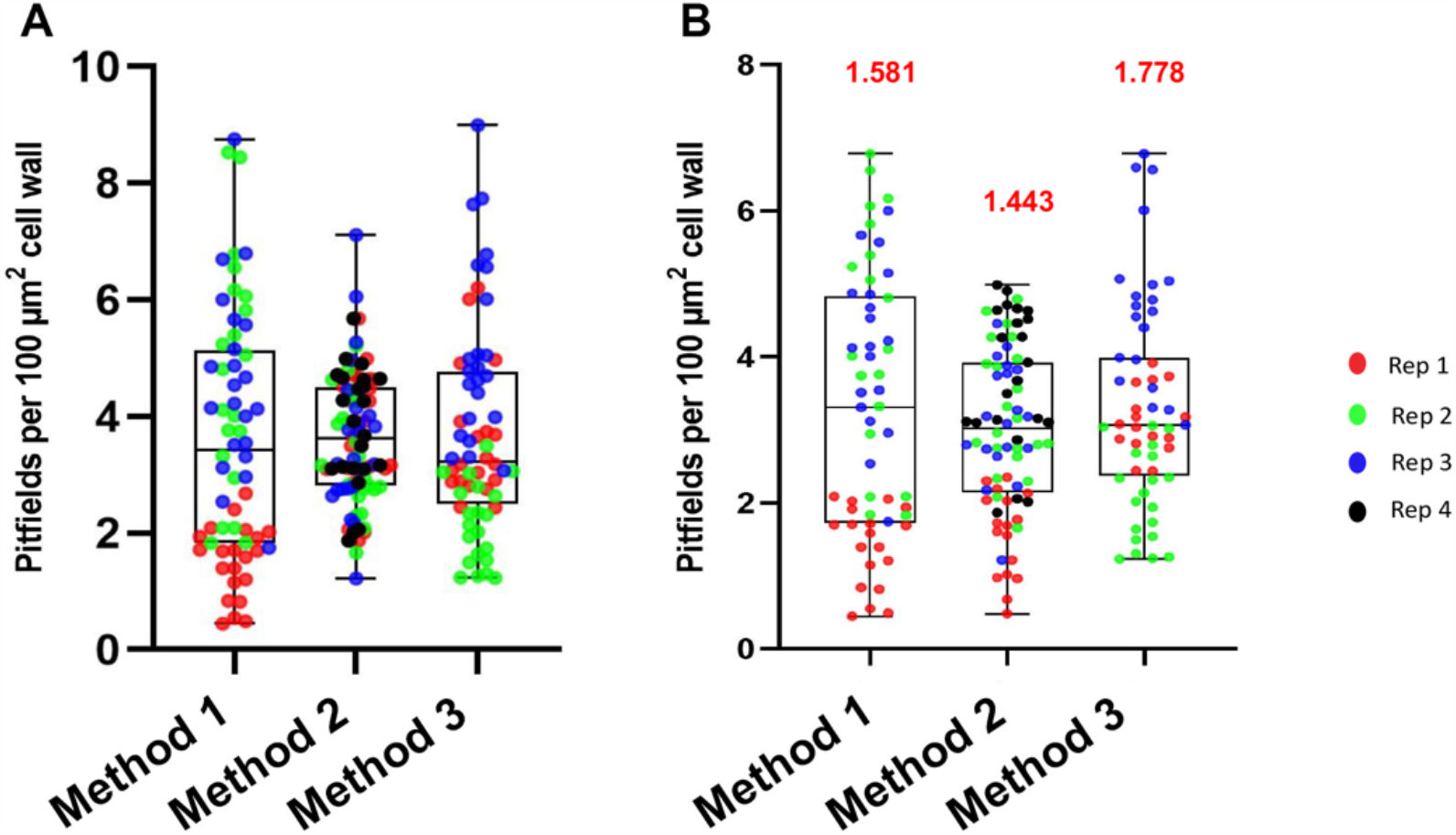
Reproducibility of pitfield quantification across methods. **A**, The pitfield quantification data from 3-4 biological replicates are shown by box and whisker plot. **B**, The same data shown in **A** is replotted using IQR analysis to remove outliers from each replicate. Median values of pitfields per 100 μm^2^ of cell wall are displayed above each plot. Medians from all methods are not significantly different from each other based on the Tukey’s multiple comparisons test. P-values for method 1 vs. method 2 combination = 0.5037; method 1 vs. method 3 = 0.9915 and method 2 vs method 3 = 0.4271.

**Figure 3.**
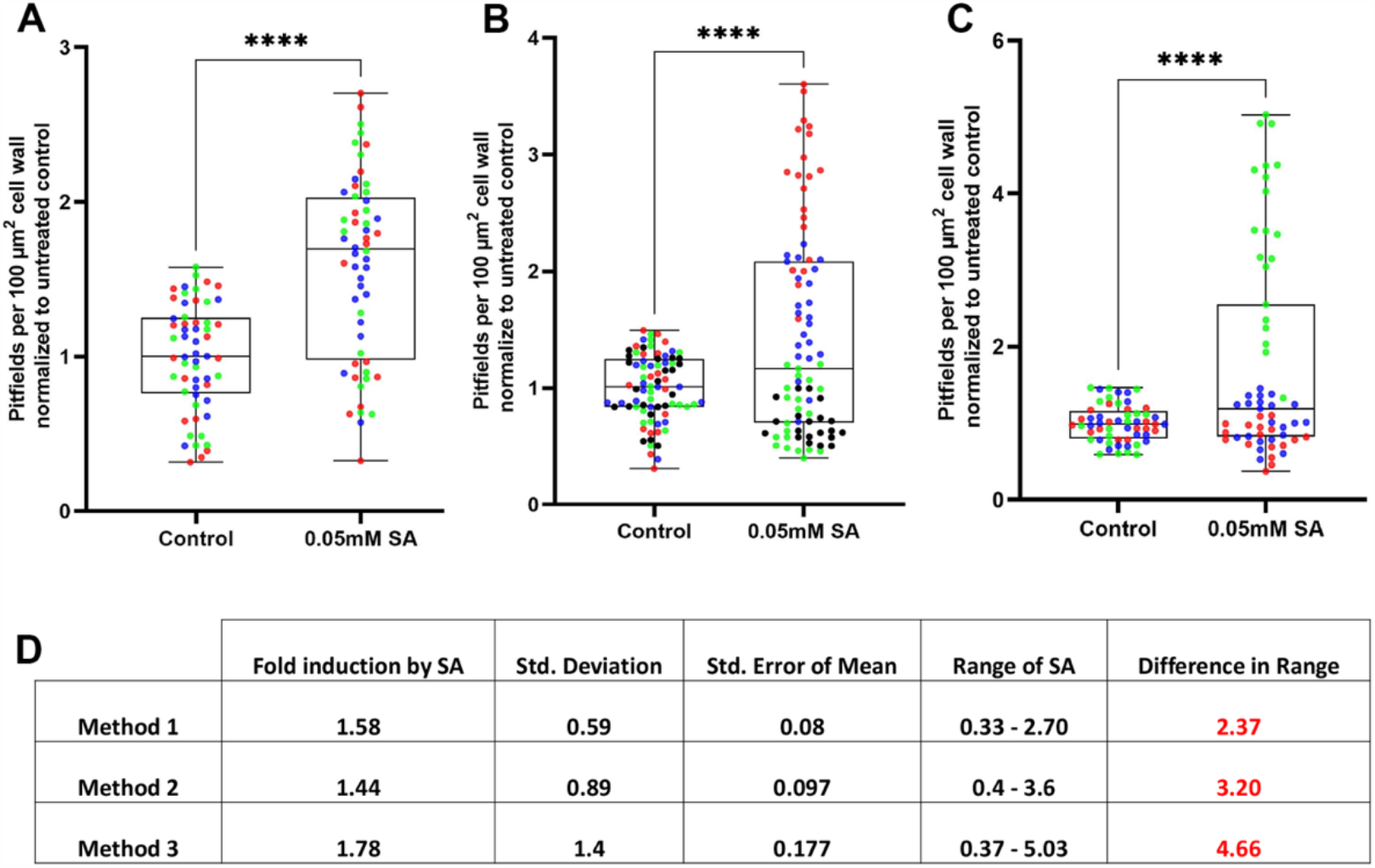
Quantification of PD-associated callose in *N. benthamiana* leaves treated with 0.05 mM salicylic acid (SA) to induce callose. Pitfield densities were quantified for at least 23 regions of interest (ROIs) for each replicate and each replicate is represented by a different color. **A**, 3 biological replicates prepared by Method 1 after 12 hours SA treatment. **B**, 4 biological replicates prepared by Method 2 after 12 hours SA treatment. **C**, 3 biological replicates prepared by Method 3 after 12 hours SA treatment. Statistical significance was determined by the Welch’s t-test (**** means p<0.0001). **D**, Fold induction by SA for all three methods along with the standard deviation, standard error and range. Pitfields per 100 μm^2^ cell wall were normalized to untreated control. An unequal variances Welch’s t-test was conducted for each plot.

A comparison of multiple pitfield density calculations from all replicates for the three methods is shown in Fig. 3. For the sake of comparison, we normalized the SA treatment data to the average control group within each replicate, and the fold induction of callose, the range, and standard deviation were determined (Fig. 3D). Comparison of the data show that Method 1 had the smallest range of values (0.33 to 2.7), with a lower occurrence of outliers and the lowest standard deviation (0.08). Data points from each replicate are distributed evenly for Method 1, whereas data points for each replicate in Methods 2 and 3 are more widely distributed over a greater range with more outliers (Fig. 3B-D). Taken together, the data show that Method 1 generates more consistent experimental results for each biological replicate with less outliers compared to the higher variability observed with Methods 2 and 3.

The generation of the data shown in Fig. 3 relies upon manual image analysis using ImageJ which is labor intensive. To evaluate whether the process could be automated, we compared the results from the manual workflow to CalloseQuant, a plug- in for semi-automated image analysis for image and quantification of callose at PD (Huang, 2022). The CalloseQuant plug-in offers two methods; CalloseQuant A is highly sensitive to fluorescence intensity whereas CalloseQuant B has low sensitivity because of thresholding, which makes it especially susceptible to the presence of fluorescence bodies (organelles) outside the ROI and outside the cell wall area (Huang, 2022). Here, both approaches were compared with the callosequant.ijm plugin for ImageJ to measure the pitfields and callose quantity of confocal images of Method 1 (Fig. 4).

**Figure 4.**
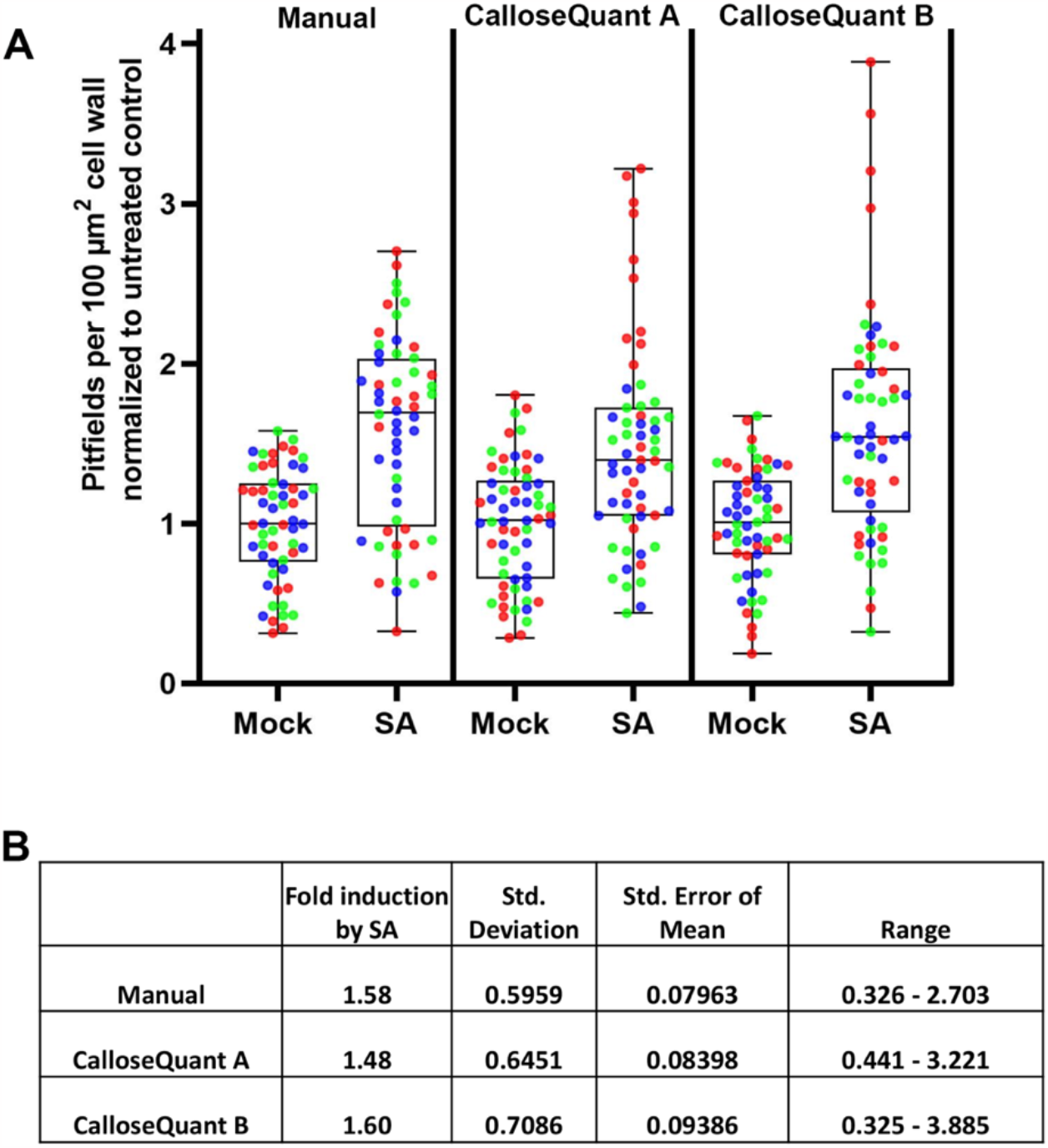
Comparison of manual and automated (CalloseQuant) methods for quantifying callose PD accumulation. **A**, Aniline blue-stained images of untreated or SA-treated samples prepared using Method 1 (fixation) were quantified using manual counting in ImageJ, CalloseQuant A, or CalloseQuant B. Three biological replicates are shown using different colors. **B**, The fold induction of callose in response to SA treatment, standard deviation, standard error and range are shown.

Images captured using confocal microscopy were first auto-scaled and converted into .tiff format using FIJI ImageJ. For the series of confocal micrographs analyzed, the peak prominence values ranges were between 25-45 μm, and the measurement radius was maintained at 5.5 for CalloseQuant A. For CalloseQuant B, the rolling ball radius range was used with 6-8 pixels, and the mean filter radius was kept at 2 pixels. We optimized other parameters described in (Huang et al., 2022) to ensure only PD-associated aniline blue-labeled foci were included in the analysis. All the data collected were manually checked for the elimination of signals and data points outside of the region of interest (ROI). A comparison of pitfield densities calculated from manual and semi-automated workflows (CalloseQuant) is shown in Fig. 4. All three analytical approaches generate datasets that show a statistically significant induction of PD callose by SA treatment. However, the median value obtained with the semiautomated approaches were lower than the manual approach and showed a wider distribution and range of data with outliers. 2D images with lower magnification are recommended for use with CalloseQuant (Huang et al., 2022). However, z-stacks (3D) images like those used in the manual workflow used in this analysis give more details on pitfields in the z-axis. When z-stacks are combined with binary watershed segmentation (which separates closely adjacent foci) a better estimation of the number of aniline blue-stained foci is likely to be obtained.

The side-by-side comparison of three methods for staining tissues for the presence of callose at PD found that fixation of *N. benthamiana* leaf tissue prior to aniline blue staining produced the most reproducible results with similar increases in callose levels on treatment with SA, (Fig. 3A and D). A comparison of manual and semi-automated analytic workflows for quantification of aniline blue-stained foci in confocal images revealed similar performance but with manual analysis a giving smaller distribution of PD density values. For high-throughput studies, the semi-automated PD quantification tool will clearly benefit the field. Recently, the use of Ilastik supervised machine learning imagery data collection software for measuring callose levels in the phloem has been reported (Welker and Levy, 2022). This approach was shown to be more reliable than the approaches using ImageJ Fiji than the manual analysis used here and CalloseQuant workflows employ. In the future, use of new analytical pipelines like those of Welker and Levy (Welker and Levy, 2022) will undoubtedly improve the quality of data regarding callose accumulation during defense.

## Acknowledgements

We thank Dr. Kirk Czymmek and staff of the Donald Danforth Plant Science Center Advanced Bioimaging Lab for assistance with microscopy and Dr. Brandon Reagan, Michigan State University, for guidance on image analysis. This work was supported by NSF award MCB 221027 to T.B.-S. and an NIH pre-doctoral fellowship 1F31GM148051- 01 to A.F.S.

---

**Fig S1.**
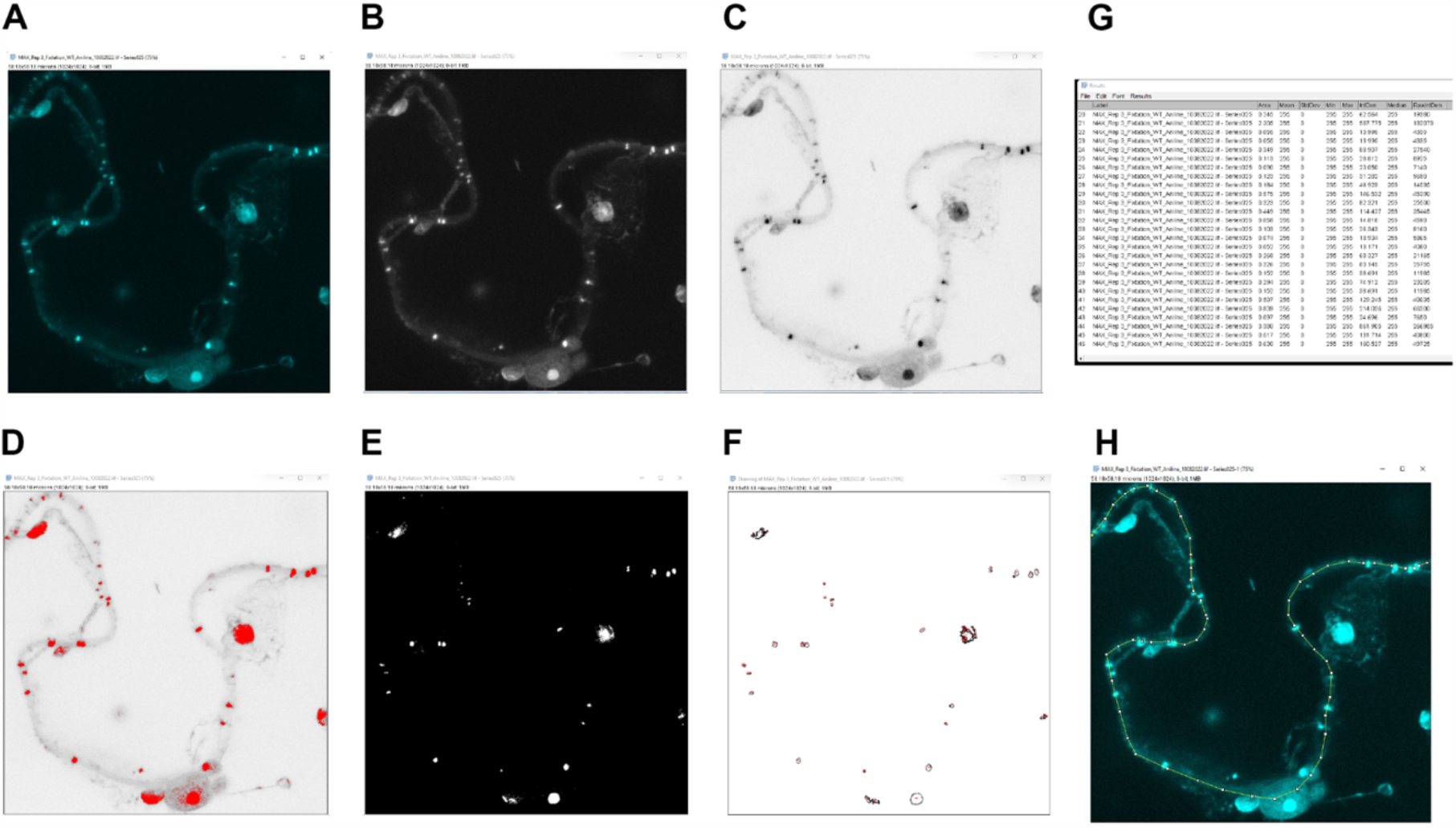
Using FIJI to manually quantify the number of pitfields and measure the cell length (area). Maximum intensity projections of a Z-stack of images (A) are converted to 8 bit (B), inverted (C), and a threshold is set to select callose staining (red; D) along the cell wall. Adjoining foci are separated to distinct bodies using binary watershed (E). Any signal that is not along the cell wall is manually excluded (F, G). Lastly, the length of cell wall within ROI is measured using the segmented line tool (H). An automatic particle analysis was performed to obtain information on PD-associated callose particles. Pitfields per 100 μm2 cell wall were calculated using Microsoft Excel.

**Fig S2.**
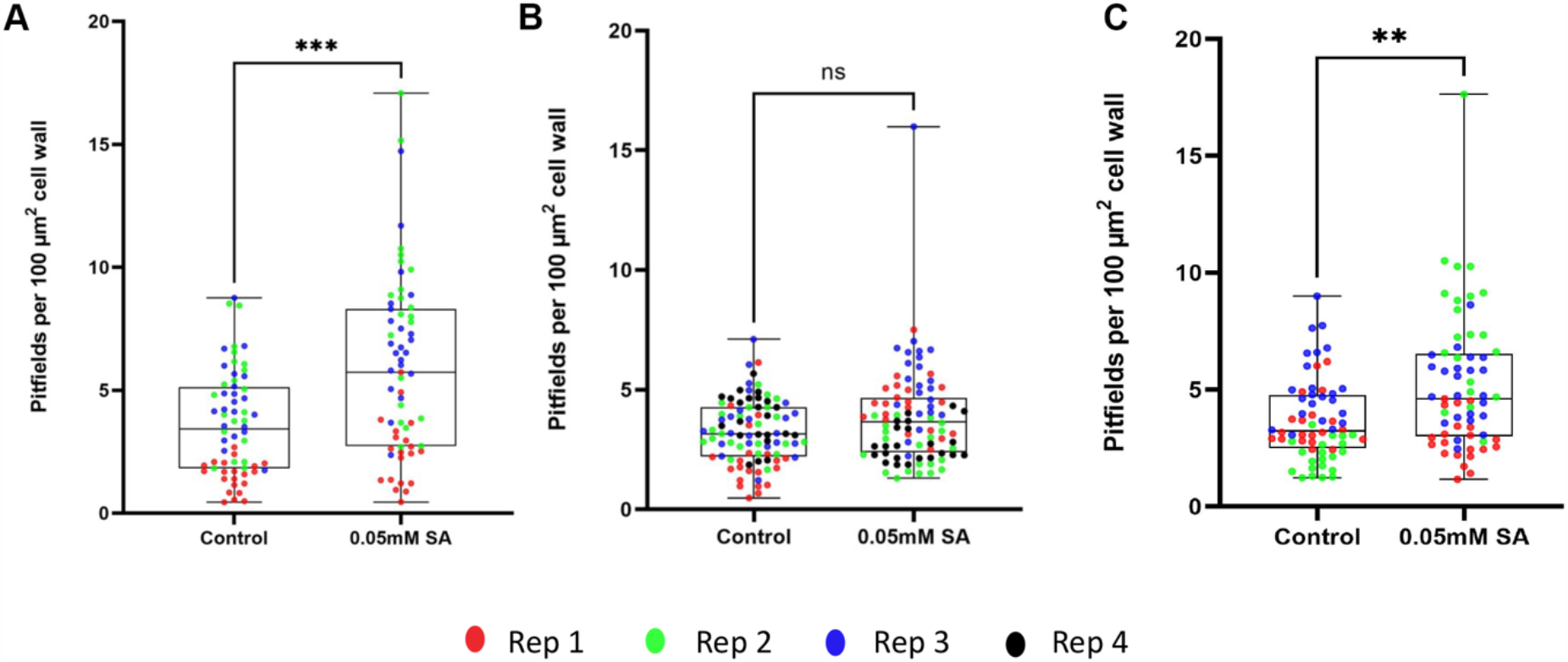
Pitfield quantifications for each method using all data, without IQR analysis. Quantification of staining of callose depositions with 1% aniline blue in untreated leaves and after 12 hours salicylic acid (SA) treatment. A, Method 1 shows a significant difference (p=0.0001), B, Method 2 show no significant difference (p=0.0705), and C, Method 3 shows a significant difference (p=0.0014) using the Welch’s t-test (ns = not significant, ** = p<0.01; ***, p<0.001).

